# Preserved Spontaneous Interpersonal Entrainment during Rhythmic Synchronization in Autism Spectrum Disorder

**DOI:** 10.64898/2026.07.09.737537

**Authors:** Maude Denis, Mattia Rosso, David Da Fonseca, Daniele Schön

**Author notes:** Correspondence: Maude Denis **Email:**.

## Abstract

**Purpose:** Interpersonal entrainment, defined as the tendency of interacting individuals to temporally align their behaviours, is considered a key mechanism supporting social interactions through predictive and multisensory processes. Autism Spectrum Disorder (ASD), characterized by social and communication difficulties, has been associated with atypical predictive and multisensory integration processes, which may affect spontaneous entrainment to other’s actions during rhythmic interactions.

**Methods:** The present study investigated spontaneous interpersonal entrainment in 24 autistic and 22 neurotypical young adults using a unidirectional adaptation of the drifting metronomes paradigm developed by Rosso et al. (2021). Participants synchronized their finger tapping with an auditory metronome while either seeing or not seeing a partner’s hand movements performing the same task at a slightly different tempo. Individual synchronization performance was assessed using asynchrony measures and computational modelling of sensorimotor synchronization, while interpersonal coordination dynamics were quantified using joint recurrence analysis.

**Results:** Visual exposure to the partner’s hand movements significantly increased tapping variability while simultaneously enhancing spontaneous interpersonal entrainment. Contrary to previous findings, these effects were comparable across groups, suggesting similar sensitivity to spontaneous low-level interpersonal coupling under stable and predictable conditions.

**Conclusion:** Overall, the findings support accounts proposing selective rather than generalized atypicalities in predictive processing and interpersonal entrainment in ASD.

## Introduction

Interpersonal entrainment (Clayton et al., 2020a; Phillips-Silver et al., 2010), defined as the tendency of interacting individuals to temporally align their behaviors, is considered as a key mechanism supporting successful social interactions. Such coordination can be observed across a wide range of contexts, for example during walking, dancing (Miyata et al., 2017; Nessler & Gilliland, 2010; Van Ulzen et al., 2008), synchronized applause (Néda et al., 2000), joint music performance (P. Keller, 2014; Schmidt & Richardson, 2008), and also conversational interactions, where interlocutors tend to align linguistic features at several levels (lexical, syntactic, semantic and prosodic) (Garrod & Pickering, 2004; Levinson & Torreira, 2015; Pickering & Garrod, 2013). Within predictive coding frameworks (Friston, 2005, 2010; Koban et al., 2019), these entrainment processes are thought to facilitate mutual prediction and reduce processing demands, thereby supporting rapid adaptation and interactional fluidity. Effective social interactions also critically rely on multisensory integration, as everyday interactions require the continuous combination of auditory and visual information. During conversation, for instance, visual cues such as lip movements, facial expressions, and gestures complement speech signals and facilitate communication (Holler & Levinson, 2019; Peelle & Sommers, 2015; Sumby & Pollack, 1954).

Because successful interpersonal coordination critically relies on multisensory integration and predictive processing, atypical functioning in these domains may influence the emergence of spontaneous interpersonal entrainment. Autism Spectrum Disorder (ASD) is a neurodevelopmental condition characterized by persistent difficulties in social communication, alongside specific and repetitive patterns of behaviour, interests, or activities, as outlined in the DSM-5 (American Psychiatric Association, 2013). Interestingly, a growing body of literature suggests that ASD may be associated with atypical sensorimotor coordination and multisensory integration processes, two mechanisms thought to play a crucial role in social interaction. In rhythmic synchronization tasks, autistic individuals have frequently been reported to show increased tapping variability (Cannon et al., 2024; Franich et al., 2021; Kasten et al., 2023; Morimoto et al., 2018; Vishne et al., 2021), reduced error correction (Kasten et al., 2023; Vishne et al., 2021) and slower adjustment to tempo perturbations (Cannon et al., 2024; Kasten et al., 2023; Vishne et al., 2021), although some contradictory findings have also been reported (Cannon et al., 2024; Sheridan & McAuley, 1997). Similarly, atypical interpersonal synchronization has been reported in ASD during both intentional rhythmic coordination tasks (Fitzpatrick et al., 2017; Kaur et al., 2018; Kawasaki et al., 2017) and spontaneous forms of entrainment (Fitzpatrick et al., 2016; Marsh et al., 2013; Trevisan et al., 2021; see also Glass & Yuill, 2024; McNaughton & Redcay, 2020 for reviews). These findings are broadly consistent with predictive coding accounts of ASD, which posits atypicalities in the formation, weighting or updating of predictions (Lawson et al., 2014; Lieder et al., 2019; Palmer et al., 2017; Sinha et al., 2014; Van de Cruys et al., 2014). In parallel, numerous studies have documented atypical multisensory integration in ASD, including lower multisensory facilitation (Brandwein et al., 2013; Collignon et al., 2013; Kwakye et al., 2011), atypical susceptibility to audiovisual illusions such as the sound-induced flash illusion (Foss-Feig et al., 2010; Kawakami et al., 2020; Stevenson, Siemann, Woynaroski, et al., 2014) and the McGurk effect (DePape et al., 2012; Mongillo et al., 2008; Stevenson, Siemann, Schneider, et al., 2014), wider temporal binding windows in simultaneity judgment tasks (Regener et al., 2024; Righi et al., 2018; E. Smith et al., 2017), and fewer benefits from visual cues during speech perception in noise (Foxe et al., 2015; E. G. Smith & Bennetto, 2007; Stevenson et al., 2017, 2018), although findings remain mixed in some paradigms (Stewart et al., 2016; Weiland et al., 2023; Woynaroski et al., 2013).

Together, these findings raise the possibility that atypical predictive and multisensory processes in ASD could affect the extent to which individuals spontaneously entrain to the actions of others during rhythmic interactions. To investigate spontaneous interpersonal entrainment in ASD, we adapted the drifting metronomes paradigm developed by Rosso et al., (2021) (see also Rosso et al., 2023, 2024). In the original version of this paradigm, participants synchronize their finger tapping with an auditory metronome while being either visually exposed or not exposed to the partner’s hand movements performing the same task at a slightly different tempo. Here, we implemented a unidirectional version of the paradigm where participants are visually exposed to the experimenter’s hand movements, while the experimenter cannot see the participant’s hand (see Figure 1). Because the experimenter’s movements are task-irrelevant and no intentional coordination is required, the paradigm allows the assessment of spontaneous interpersonal coupling emerging from low-level sensorimotor interactions. The present study examined whether participants with ASD differ from neurotypical participants in the extent to which visual interpersonal information influences individual synchronization performance and dyadic coordination dynamics. Based on previous findings of atypical sensorimotor synchronization and multisensory integration in ASD, we expected reduced spontaneous interpersonal entrainment in autistic participants, reflected by weaker coupling-related changes in synchronization behavior and dyadic coordination dynamics.

**Fig. 1.**
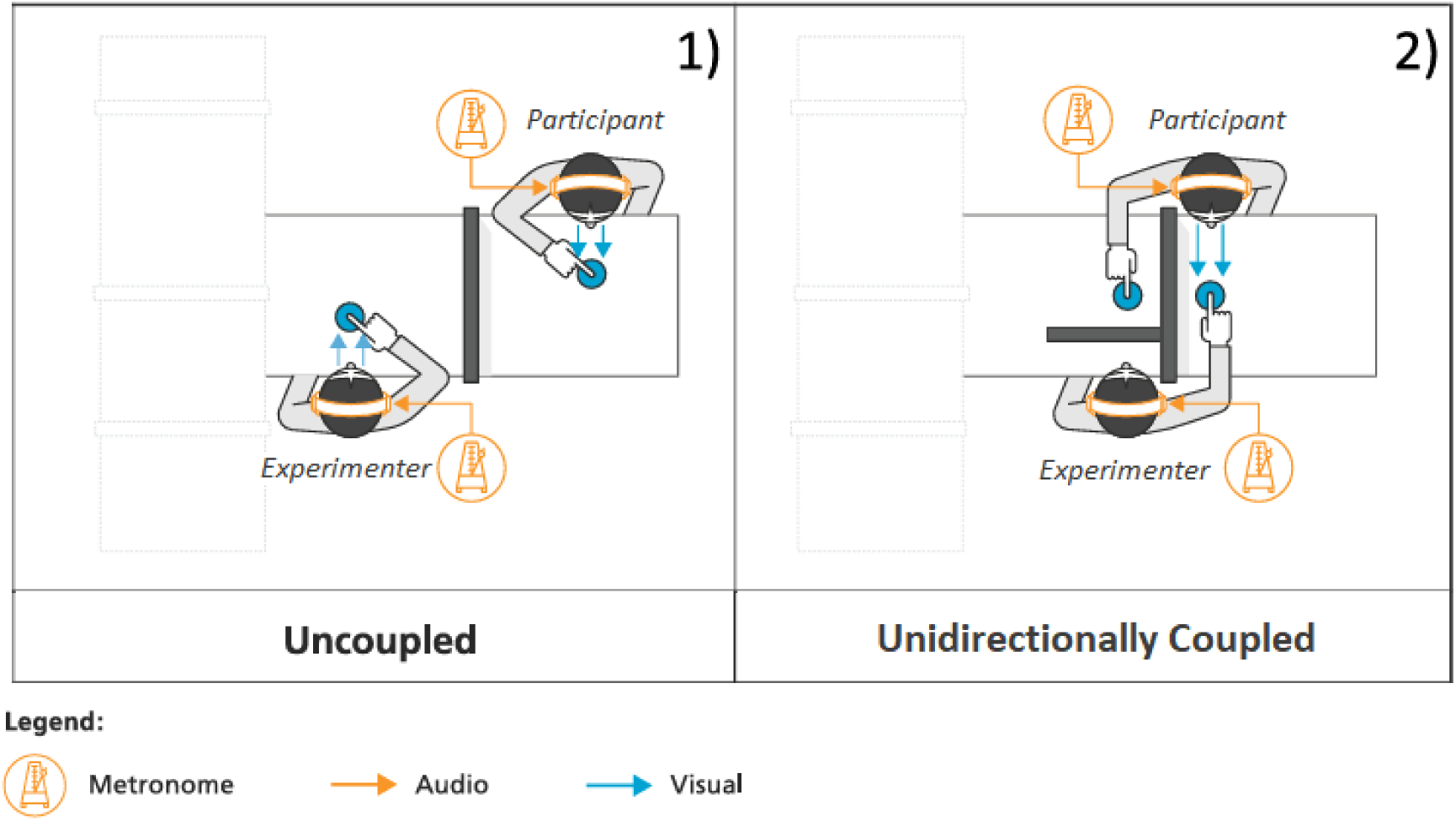
Experimental design. Participants performed a finger-tapping task while synchronizing with an auditory metronome in two conditions. In the *uncoupled condition* (baseline), participants synchronized their tapping with their own metronome while visually focusing on their own hand. The view of the experimenter’s hand was hidden by a screen placed on the table. In the *unidirectional coupled condition*, participants synchronized with their own metronome while looking at the experimenter’s hand tapping at a slightly different tempo and were instructed to ignore these movements and focus on their own metronome. The experimenter could not see the participant’s hand in either condition, ensuring a unidirectional visual influence from the experimenter to the participant

## Materials and Methods

### Participants

Twenty-four young adults with autism spectrum disorder (ASD; 18 - 36 years) and twenty-two neurotypical controls participated in the study. Participants were recruited through advertisements in the Developmental Psychiatry Unit of the Salvator Hospital (AP-HM), through two independent patient associations located in Marseille (France), and through Aix-Marseille University (France). ASD diagnoses were established by experienced clinicians according to DSM-5 criteria, supported when available by standardized diagnostic tools (e.g., ADOS-2, ADI-R).

Groups were comparable in age (t = −1.20, p = 0.24; ASD: M = 24.2 y, SD = 4.49; Controls: M = 22.9 y, SD = 2.97) and years of education (t = 1.50, p = 0.15; ASD: M = 3.4 y, SD = 1.90; Controls: M = 4.4 y, SD = 1.61), and showed no statistically significant difference in gender distribution (χ² = 2.49, p = 0.11; ASD: 12F/12M; controls: 6F/16M).

Exclusion criteria included neurological, sensory, motor, or language disorders in all participants, and ASD or other neurodevelopmental disorders in the control group. Additional exclusion criteria for the ASD group comprised psychotic disorders, known genetic or medical conditions associated with autism, severe perinatal complications and central nervous system infections.

All participants provided written informed consent. The experimental protocol was evaluated by the Aix-Marseille Ethics committee and all procedures complied with the Declaration of Helsinki. Participants could opt out of the study at any time and were compensated for their participation.

### Procedure

The paradigm was adapted from Rosso et al. (2021). The participant and the experimenter sat across a table, facing each other. Each tapped with their dominant hand on an individual wooden box containing a microphone (DPA 4099), connected to a Scarlett 2i2 soundcard (24-bit, 192 kHz).

The participant and the experimenter were presented with slightly different auditory metronome tempi (respectively 1.67 Hz and 1.64 Hz). The metronome signals were initially aligned at 0° relative phase, which then progressively shifted in regular 5.6° steps, resulting in a complete cycle every 39 s. Five consecutive cycles were performed in each experimental condition. Auditory stimuli were delivered through headphones (t.bone HD 1800) via independent audio channels. Volume was individually adjusted before the experiment, and pink noise was used to mask external sounds.

In the uncoupled tapping (baseline) condition, the participant and the experimenter synchronized with their respective metronome while looking at their own hand, without seeing or hearing each other (see Figure 1.1).

In the unidirectional visual coupling condition, participants were instructed to look at the experimenter’s hand but to ignore it and continue tapping along with their assigned metronome. The experimenter could not see the participant’s hand and focused on her own tapping throughout the task to minimize potential bias arising from her awareness of participants’ diagnostic status, as well as potential learning effects across participants, thereby implementing a scenario of unidirectional visual coupling (see Figure 1.2).

### Analysis

Individual synchronization performance was quantified using asynchronies, defined as the time interval between each metronome event and the corresponding participant’s tap. Mean and standard deviation of asynchronies were used as measures of tapping accuracy and variability. In addition, tapping consistency was assessed using circular statistics, with R corresponding to the length of the mean vector (ranging from 0 to 1). Higher R values indicate more consistent tap timing relative to the metronome phase. As a second approach, we applied a computational model of sensorimotor synchronization (Vorberg & Schulze, 2002; Vorberg & Wing, 1996; Wing & Kristofferson, 1973). In this model, the interval between successive taps is assumed to be influenced by three components: timekeeping of the metronome tempo, motor execution, and phase correction (α), defined as the proportion of the previous asynchrony that is corrected in the next tap. Higher α value indicates stronger phase correction. Model parameters were estimated using the bGLS method introduced by Jacoby et al. (2015) and adapted by Vishne et al (2021) to accommodate missing data.

Unidirectional coupling dynamics were assessed using joint recurrence analysis, following the framework proposed by Rosso et al. (2021). Cleaned time series were interpolated using a sine function (1 kHz sampling rate) and downsampled by a factor of 40 prior to analysis. Time-delay embedding parameters were optimized for each participant and average parameter values were subsequently applied across datasets. Individual recurrence plots (RPs) were computed for each participant. Joint recurrence plots (JRPs) were then obtained for each dyad by overlapping both individual RPs and retaining only simultaneous recurrences. The five trials of each condition were aggregated, yielding a one-dimensional recurrence score reflecting the density of coupled behavior across the metronome cycle (for a detailed methodological description, see Rosso et al., 2021).

### Statistical analysis

All statistical analysis were conducted in R (R Core Team, 2024) using the lme4 (Bates et al., 2015) for mixed-effects modelling and emmeans packages (Lenth et al., 2022), for post-hoc comparisons.

Individual synchronisation performance was assessed using linear mixed-effects models with group and coupling condition as fixed effect and participant as a random effect. Dependent variables (denoted here as x) included negative mean asynchrony (NMA), asynchrony variability (SD), tapping consistency (R), timekeeper noise, motor noise, and phase correction (α). For each variable, the main model was compared with a null model (lmer(x ∼ 1 +(1|participant)) and with a model including coupling condition and its interaction with group (lmer(x ∼ group * condition + (1|participant)). Due to non-normality distribution, tapping consistency (R) was analysed separately using Wilcoxon rank-sum tests.

Recurrence scores were analysed using mixed-effects models including group and coupling condition as fixed factors and Time as a continuous predictor corresponding to the 64 metronome phase steps. Linear and quadratic orthogonal polynomials of Time were included to characterize temporal changes in coupling dynamics. Dyad-specific random effects were included to account for inter-dyad variability in synchronization and coupling susceptibility. The formula of the full model is the following:

Recurrence ∼ (Time +Time^2^) ∗ Group ∗Coupling + (Time +Time^2^ | Dyad) + (Time+Time^2^ | Dyad : Group : Coupling)

Model comparison was used to evaluate the statistical significance of fixed effects, based on the Akaike Information Criterion (AIC), which provides a balance between model complexity and explanatory power. Residual normality and homoscedasticity were visually inspected for all models. Post-hoc contrasts were computed using emmeans with Tukey correction for multiple comparisons.

Complementary Bayesian mixed-effects analyses were conducted using the brms package in R (Bürkner, 2017). Bayesian models mirrored the structure of the frequentist mixed-effects models and were estimated using Hamiltonian Monte Carlo sampling implemented in Stan. Weakly informative priors were used for fixed and random effects parameters. Posterior distributions were summarized using posterior means and 95% credible intervals (CrIs), and convergence was assessed using Rhat statistics and effective sample size estimates.

### Results Individual synchronization performance

Negative mean asynchrony (NMA) did not significantly differ between groups (β = 0.002, χ² = 0.06, p = 0.80), between coupling conditions (β = 0.007, χ² = 0.91, p = 0.34), and no Group x Coupling interaction was observed (β = −0.009, χ² = 0.36, p = 0.55), indicating comparable synchronization accuracy across groups and conditions.

In contrast, the coupling condition significantly affected synchronization stability. The standard deviation of asynchrony (SD) was significantly higher in the unidirectional coupled condition than in the uncoupled condition (β = −0.032, χ² = 14.98, p < 0.001; Coupled: M = 0.09, SD = 0.038; Uncoupled: M = 0.06, SD = 0.041; see Figure S1. A.) indicating greater temporal variability when participants were visually exposed to the experimenter’s movements. Similarly, tapping consistency (R) was significantly lower in the coupled condition (W = 407, p < 0.001, r = 0.544; Coupled: M = 0.70, SD = 0.18; Uncoupled: M = 0.86, SD = 0.15; see Figure S1. B.). These effects did not differ between groups (Group x Coupling; SD: β = 0.009, χ² = 0.31, p = 0.58; R: W = 332, p = 0.24, r = 0.174).

Computational modelling of sensorimotor synchronization further supported these findings. Phase correction, reflecting the proportion of the previous asynchrony corrected on the subsequent tap, was significantly reduced in the unidirectional coupled condition relative to the uncoupled condition (β = 0.180, χ² = 24.24, p < 0.001; coupled: M = 0.13, SD = 0.13; uncoupled: M = 0.31, SD = 0.20, see Figure S1. C.). This modulation of phase correction was similar across groups (Group x Coupling: β = −0.016, χ² = 0.06, p = 0.81).

We next examined whether these alterations in individual synchronization behavior were associated with changes in dyadic coordination dynamics.

### Dyadic coordination dynamics

Joint recurrence analysis revealed a significant main effect of coupling condition on dyadic coordination dynamics. Recurrence scores, indexing the degree of spontaneous interpersonal entrainment, were significantly higher in the unidirectional coupled condition compared to the uncoupled condition (β = 9.526, t = 4.19, p < 0.001; Coupled: M = 68.3, SD = 20.3; Uncoupled: M = 59.2, SD = 11.3). In addition, significant linear and quadratic Time × Coupling interactions were observed (linear: β = −44.21, t = −4.39, p < 0.001; quadratic: β = 30.50, t = 2.40, p = 0.016), indicating that the temporal profile of dyadic recurrence differed across the metronome cycle in the coupled condition, with stronger entrainment around specific phase relations, particularly near in-phase attractor regions (see Figure 2).

**Fig. 2.**
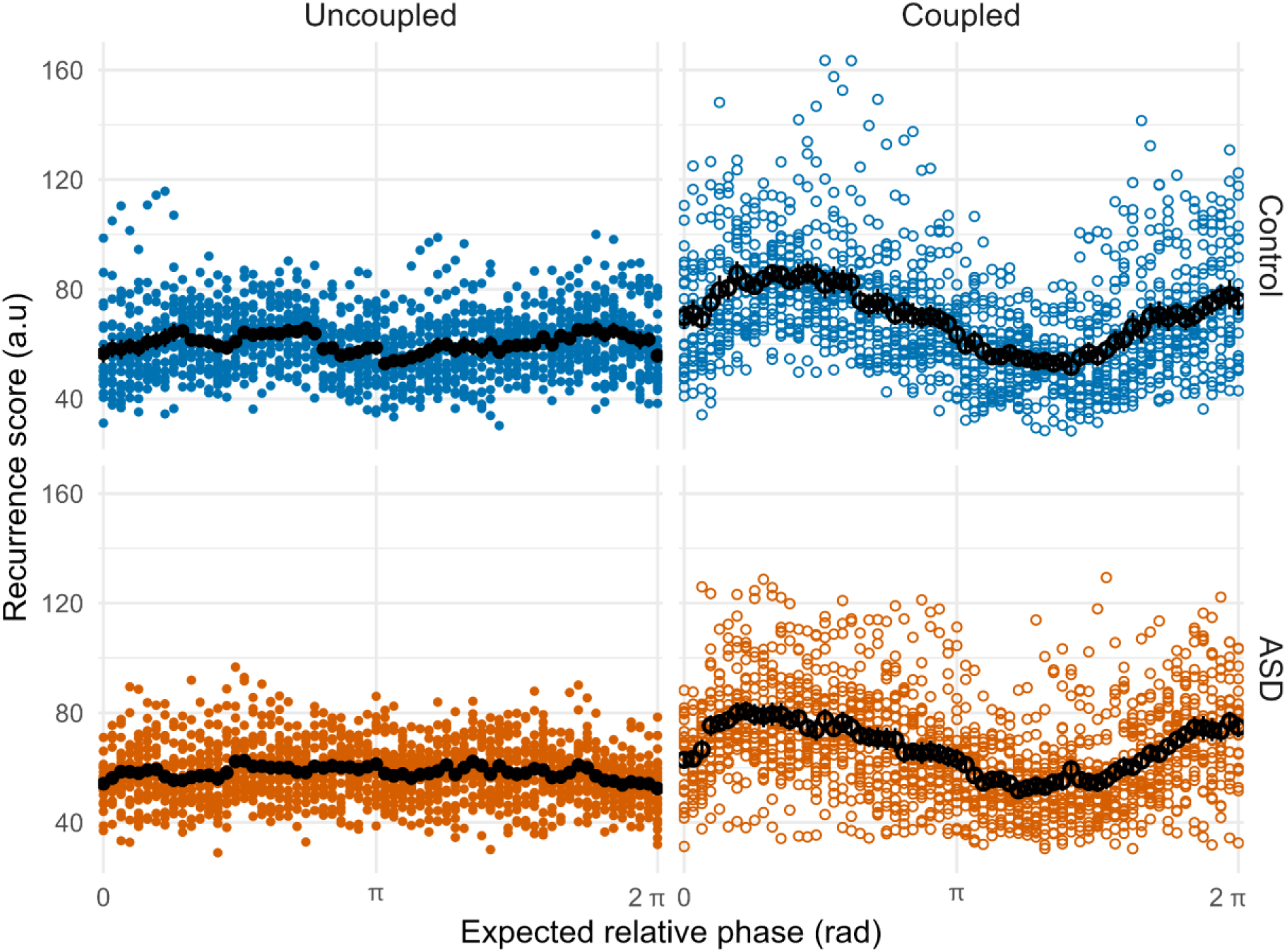
Dyadic coordination dynamics. Joint recurrence analysis was used to quantify spontaneous interpersonal entrainment across the metronome phase cycle. Visual coupling increased overall recurrence relative to the uncoupled condition and induced phase-dependent modulations of dyadic coordination, with stronger entrainment around in-phase regions. Effects were comparable between groups. Grand-average and error bars are represented in black

Crucially, this enhancement of dyadic entrainment did not differ between groups (Group x Coupling: β = −0.789, t = - 0.25, p = 0.80), suggesting comparable sensitivity to spontaneous interpersonal coupling in autistic and neurotypical participants.

Complementary Bayesian mixed-effects modelling yielded similar conclusions. Visual coupling was associated both with higher overall recurrence and with stronger phase-dependent fluctuations in recurrence across the metronome cycle, particularly around in-phase regions. In contrast, posterior estimates for the Group × Coupling interaction were centered near zero (β = −0.00, 95% CrI [-0.37, 0.37]), providing little evidence for group-related differences in coupling effects. Likewise, higher-order interactions involving Group and the temporal components of the metronome cycle broadly overlapped zero (see Table S1).

## Discussion

The present study investigated whether spontaneous visual interpersonal coupling during rhythmic synchronization differs in autistic and neurotypical young adults. Results showed that visual exposure to the experimenter’s hand movements influenced participants’ synchronization behavior despite explicit instructions to ignore these movements. More specifically, visual coupling increased temporal variability in tapping while simultaneously enhancing spontaneous dyadic entrainment, particularly around in-phase attractor regions, consistent with previous findings in neurotypical populations (Rosso et al., 2021, 2023, 2024). Crucially, these effects were comparable across groups, suggesting similar sensitivity to spontaneous low-level interpersonal coupling under the present experimental conditions.

The absence of group differences contrasts with previous reports of atypical sensorimotor synchronization, multisensory integration, and interpersonal coordination in ASD (see Denis et al., 2025; McNaughton & Redcay, 2020; Stevenson et al., 2016 for reviews). Prior studies have frequently reported increased tapping variability and reduced phase correction during rhythmic synchronization tasks, as well as atypical audiovisual integration and atypical interpersonal coordination in both explicit and spontaneous interaction contexts. In the present study, however, visual exposure to another person’s movements influenced synchronization behavior and dyadic entrainment similarly in autistic and neurotypical participants. These findings therefore suggest that spontaneous low-level interpersonal coupling may remain preserved in ASD under certain controlled interactional conditions.

One potential explanation for the present results is that atypicalities in ASD may emerge primarily in contexts requiring rapid online adaptation and integration of complex or highly dynamic multimodal information, whereas performance may remain relatively preserved when temporal dynamics are slower and sensory input remains simple and predictable. In the present experiment, metronome dephasing unfolded over a relatively long temporal scale (39-s cycle duration) and involved relatively simple rhythmic and visual stimuli, potentially allowing participants with ASD to adapt their behaviour in a comparable manner to neurotypical participants. This interpretation is consistent with the *slow updating hypothesis* of ASD, which posits delayed updating of predictive models rather than a generalized impairment in prediction or synchronization processes (Kasten et al., 2023; Lieder et al., 2019; Vishne et al., 2021). More broadly, the present findings suggest that spontaneous interpersonal coupling mechanisms may be preserved in ASD when interaction dynamics evolve slowly and predictably, whereas difficulties may emerge under conditions requiring faster predictive updating and adaptation.

These findings may have important implications for predictive processing accounts of ASD (Lawson et al., 2014; Palmer et al., 2017; Sinha et al., 2014; Van de Cruys et al., 2014). Within predictive coding frameworks, spontaneous interpersonal entrainment has been proposed to support social interaction by minimizing mismatches between self-generated and observed actions during dyadic interaction (Koban et al., 2019; Rosso et al., 2021). The presence of spontaneous visual interpersonal coupling in both groups therefore suggests that low-level predictive mechanisms supporting interpersonal coordination may not be globally impaired in ASD. Rather, the present findings are more consistent with accounts proposing selective atypicalities in predictive processing, such as *slow updating* (Kasten et al., 2023; Lieder et al., 2019; Vishne et al., 2021) or *hypervolatility hypothesis* (Lawson et al., 2017; Palmer et al., 2017; Van de Cruys et al., 2014). More broadly, these results challenge strongly generalized interpretations of predictive atypicalities in ASD and instead support the idea that predictive and interpersonal coordination processes may be preserved under sufficiently stable and predictable interactional conditions.

Spontaneous entrainment processes contribute to many everyday social interactions, including conversational alignment (Garrod & Pickering, 2004), rhythmic coordination (Clayton et al., 2020b; P. E. Keller et al., 2014), and turn-taking (Garrod & Pickering, 2015; Levinson, 2016). The present findings therefore raise the possibility that spontaneous low-level coordination may act as a scaffold for higher-order social coordination during interpersonal interaction in ASD (Fiveash et al., 2021; Fram et al., 2024; Knoblich & Sebanz, 2008; Lense & Camarata, 2020).

Several limitations should nevertheless be acknowledged. First, the modest sample size may have reduced sensitivity to subtle group differences, particularly given the heterogeneity of ASD. Second, the use of a highly controlled rhythmic interaction limits the ecological validity of the findings compared to more naturalistic social contexts involving richer and less predictable multimodal exchanges. Third, the present paradigm specifically targeted spontaneous low-level sensorimotor coupling and therefore does not necessarily generalize to higher-order forms of interpersonal coordination involving explicit social intentions, conversational dynamics, or complex joint actions. Future studies should examine whether similar patterns are observed in more naturalistic and socially demanding interactional settings.

In summary, the present study shows that spontaneous visual interpersonal coupling during rhythmic synchronization similarly influenced autistic and neurotypical adults in a controlled drifting metronomes paradigm. Although visual coupling reduced individual synchronization stability, it simultaneously enhanced dyadic entrainment in both groups, particularly around in-phase attractor regions. These findings suggest that low-level spontaneous entrainment mechanisms may be preserved in ASD and could provide a building block for more complex forms of interpersonal coordination, especially in stable and predictable interactional contexts.

## Author Contributions

Maude Denis: Conceptualization, Methodology, Formal analysis, Project Administration, Investigation, Writing – original draft, Writing – review & editing, Visualization. Mattia Rosso: Conceptualization, Methodology, Writing – review & editing. David Da Fonseca: Writing – review & editing, Supervision. Daniele Schön: Conceptualization, Funding Acquisition, Project Administration, Writing – original draft, Writing – review & editing, Supervision.

## Acknowledgments

We thank the participants for their time and commitment, the Association Soliane, Autisme PACA and Tiffany Esnos for their support with participant recruitment, and Isabelle Charvin for her assistance with data preparation. Research was supported by grants ANR-21-CE28-0010 (DS), ANR16-CONV-0002 (ILCB), ANR-17-EURE-0029 (NeuroMarseille), and the Excellence Initiative of Aix-Marseille University (A*MIDEX).

