## Supplementary Material for "Preserved Spontaneous Interpersonal Entrainment during Rhythmic Synchronization in Autism Spectrum Disorder"

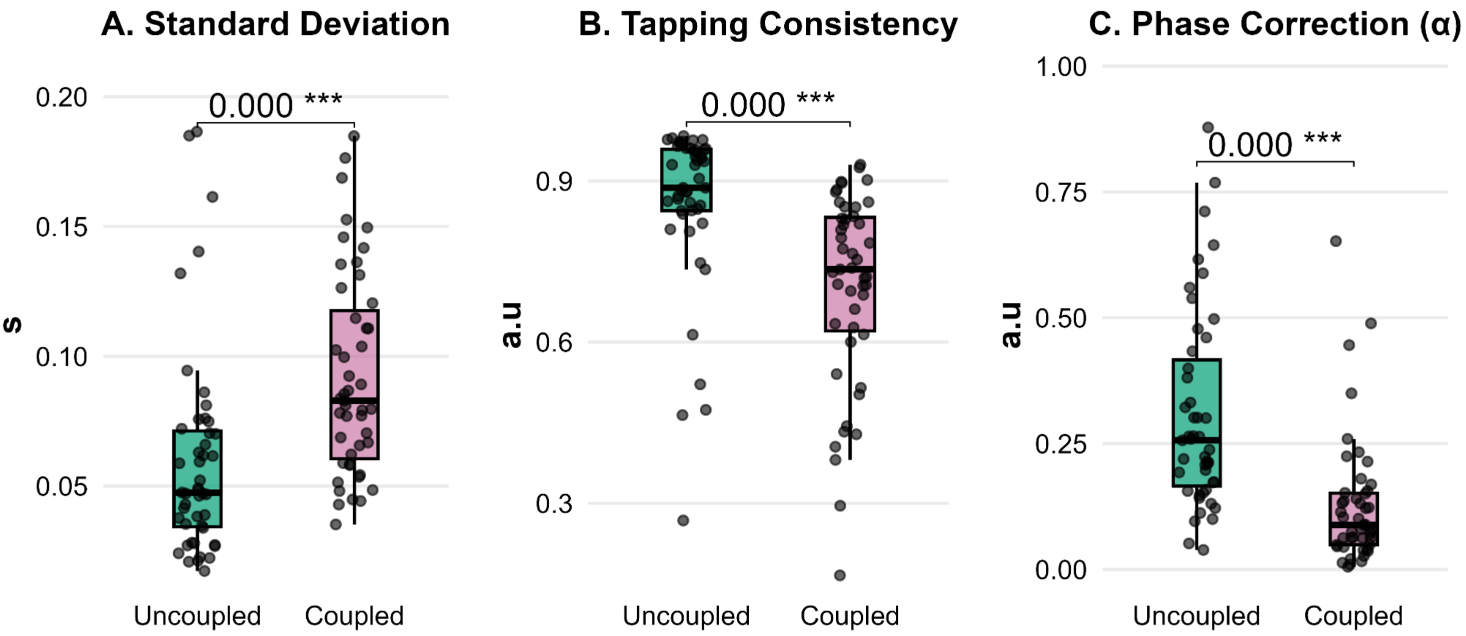


**Fig. S1 Individual synchronization performance**

(A) Standard deviation of asynchrony (SD). (B) Tapping consistency (R), higher values indicate greater consistency. (C) Phase correction parameter ($\alpha$) from computational modeling, representing the proportion of the previous asynchrony corrected on the subsequent tap. For all panels, boxplots show median, interquartile range, and individual data points. Uncoupled: green; coupled: purple. N = 24 ASD, 22 controls. Significance: $*p < 0.05$, $**p < 0.01$, $***p < 0.001$

| Effect | Estimate | 95% CrI | Interpretation |
|---|---|---|---|
| Coupling | 0.50 | [0.23 ; 0.77] | Higher recurrence in the coupled condition |
| Time (linear) × Coupling | -1.86 | [-2.74 ; -0.97] | Coupling altered the linear evolution of recurrence across the cycle |
| Time (quadratic) × Coupling | 1.56 | [0.53 ; 2.61] | Coupling altered the nonlinear recurrence profile across the cycle |
| Group × Coupling | -0.00 | [-0.37 ; 0.37] | Little evidence for group differences in coupling effects |

**Table S1 Bayesian estimates of coupling effects on dyadic recurrence dynamics**

Estimates correspond to posterior means obtained from the Bayesian mixed-effects model of recurrence dynamics. CrIs = 95% credible intervals. Time (linear) and Time (quadratic) correspond to the first- and second-order orthogonal polynomial components modelling temporal changes in recurrence across the metronome cycle. Positive estimates indicate increased recurrence scores in the coupled condition
